# Genome-wide characterization of RNA editing in chicken: lack of evidence for non-A-to-I events

**DOI:** 10.1101/008912

**Authors:** Laure Frésard, Sophie Leroux, Pierre-François Roux, Christophe Klopp, Stéphane Fabre, Diane Esquerré, Patrice Dehais, Anis Djari, David Gourichon, Sandrine Lagarrigue, Frédérique Pitel

## Abstract

RNA editing corresponds to a post-transcriptional nucleotide change in the RNA sequence, creating an alternative nucleotide, not present in the DNA sequence. This leads to a diversification of transcription products with potential functional consequences. Two nucleotide substitutions are mainly described in animals, from adenosine to inosine (A-to-I) and from cytidine to uridine (C-to-U). This phenomenon is more and more described in mammals, notably since the availability of next generation sequencing technologies allowing a whole genome screening of RNA-DNA differences. The number of studies recording RNA editing in other vertebrates like chicken are still limited. We chose to use high throughput sequencing technologies to search for RNA editing in chicken, to understand to what extent this phenomenon is conserved in vertebrates.

We performed RNA and DNA sequencing from 8 embryos. Being aware of common pitfalls inherent to sequence analyses leading to false positive discovery, we stringently filtered our datasets and found less than 40 reliable candidates. Conservation of particular sites of RNA editing was attested by the presence of 3 edited sites previously detected in mammals. We then characterized editing levels for selected candidates in several tissues and at different time points, from 4.5 days of embryonic development to adults, and observed a clear tissue-specificity and a gradual editing level increase with time.

By characterizing the RNA editing landscape in chicken, our results highlight the extent of evolutionary conservation of this phenomenon within vertebrates, and provide support of an absence of non A-to-I events from the chicken transcriptome.

## BACKGROUND

A fascinating reality of the genome, receiving more and more empirical evidences, is that its biology is far more complex than previously thought. The rule “one gene has one DNA sequence leading to one mRNA translated into one protein”, even if not (yet) an exception, is now well-known to be transgressed in a vast field of possibilities. Taking the example of the human genome, the number of genes, the percentage of the genome that is transcribed, the alternative transcripts count per gene, or the way their expression is regulated, are all characteristics for which knowledge is moving with an extraordinary pace. The ENCODE project brought a lot of data and analyses in this line [1]. Among transformations that RNA transcripts undergo during maturation, RNA editing is a phenomenon leading to differences between the final RNA sequence and the DNA region it was transcribed from. The term was first used by Benne *et al* in 1986 [2], and can now be defined, in a broad sense, as a nucleotide insertion, deletion or substitution in the RNA sequence, occurring in various types of RNA, from tRNA to mRNA, either coding or not [3]. Substitutions comprise several types of modifications, the most common in vertebrates being the A-to-I conversion, catalyzed by the ADAR family enzymes (Adenosine Desaminase that Acts on RNA) [4] and leading to an A-to-G reading of the cDNA molecule [5, 6] and C-to-U conversion, catalyzed by the APOBEC enzyme [7, 8].

RNA editing is limited to eukaryotes, with a few exceptions (see [9] for review). It is observed in chloroplasts, widespread in mitochondria, and also found as a nuclear phenomenon in animals. It seems to have arisen through different mechanisms in different lineages, rather than being inherited from a common ancestor, and whether natural selection was involved in its evolution is still debated [9–11]. While RNA editing is more and more characterized in mammals, especially in human, mouse and rat [12–18], only a few studies have been performed in birds and were targeting specific genes. The apolipoprotein B (APOB) RNA editing, well-known in mammals, seems to be absent from chicken [19] and zebra finch [20]. In chicken, the CYFIP2 (cytoplasmic FMR1 interacting protein 2) and FLNA (filamin A) genes are edited in brain and liver [21], the splicing regulator NOVA1 (Neuro-Oncological Ventral Antigen 1) is edited in the brain [22], the GABA_A_ (gamma-Aminobutyric Acid Type A) Receptor, alpha3 subunit (GABRA3) is edited in the brain and the retina [23, 24]. But no genome-wide study is available to really assess the extent of RNA editing in this species. High-throughput RNA sequencing actually allows performing a deeper transcriptome analysis than previous technologies, including RNA editing through a genome-wide approach [25]. This has been performed on several species, including human and mouse [12, 13, 15, 26, 27] but never in avian species. The number of editing sites (or detected as RDD: RNA-DNA Differences) observed in mammals strongly varies between studies, even on the same tissues of the same species, and an increasing number of analyses point the requirement of very careful bioinformatics procedures to limit technical artifacts [14, 15, 28–32].

To improve the available knowledge about the extent of RNA editing in chicken, we chose an approach without *a priori* by using DNA and RNA sequencing on the same samples through Next Generation Sequencing (NGS) technology of whole embryos. Our results support the fact that RNA editing is not a frequent event in chicken, is mostly limited to the canonical A-to-I conversions, and shows strong tissue- and developmental-specificities.

## RESULTS

### Sequences analysis

DNA and RNA sequences were obtained from the same samples of chicken embryos. In average, 141,534,451 DNA reads and 65,302,559 RNA reads were aligned and analyzed for each embryo. The genome coverage reaches 93% for DNA reads and 22% for the RNA reads. A summary on DNA and RNA sequences aligned on Galgal4 chicken assembly is presented in Table 1.

**Table 1.** Number of analyzed RNA and DNA sequences in the study (after alignment on Galgal4)

|  | Embryo (n=8) |
| --- | --- |
| Mean total number of reads (DNA) | 141 534 451 |
| Mean total number of reads (RNA) | 65 302 559 |
| Total number of reads (DNA) | 1 132 275 604 |
| Total number of reads (RNA) | 522 420 469 |
| Mean coverage (DNA) - min coverage 5 reads (% genome) | 93.1 ± 1.0 |
| Mean coverage (RNA) - min coverage 5 reads (% genome) | 22.3 ± 2.1 |
| Mean coverage (RNA) - min coverage 15 reads (% genome) | 16.4 ± 2.6 |

### Data filtering–biases detection

The first step was to detect RDD sites, *i.e.* positions homozygous in DNA and presenting an alternative sequence in RNA. To consider a position as potentially candidate, we fixed a minimum read-depth threshold of 15 both in DNA and RNA alignments for each embryo. We only kept candidates for which the alternative nucleotide frequency in DNA was null (Figure 1A). A total of 1,327 RDD sites met this criterion. The next filtering steps are aiming to avoid common pitfalls in sequences analysis, in order to decrease the number of putative false positive RDD candidates (Figure 1). To increase the robustness of the results and avoid putative false positive due to an artifact present only in one sample, we only considered RDD sites detected in at least 2 biological replicates. We ended up with 324 RDD sites (Figure 1B).

**Figure 1.**
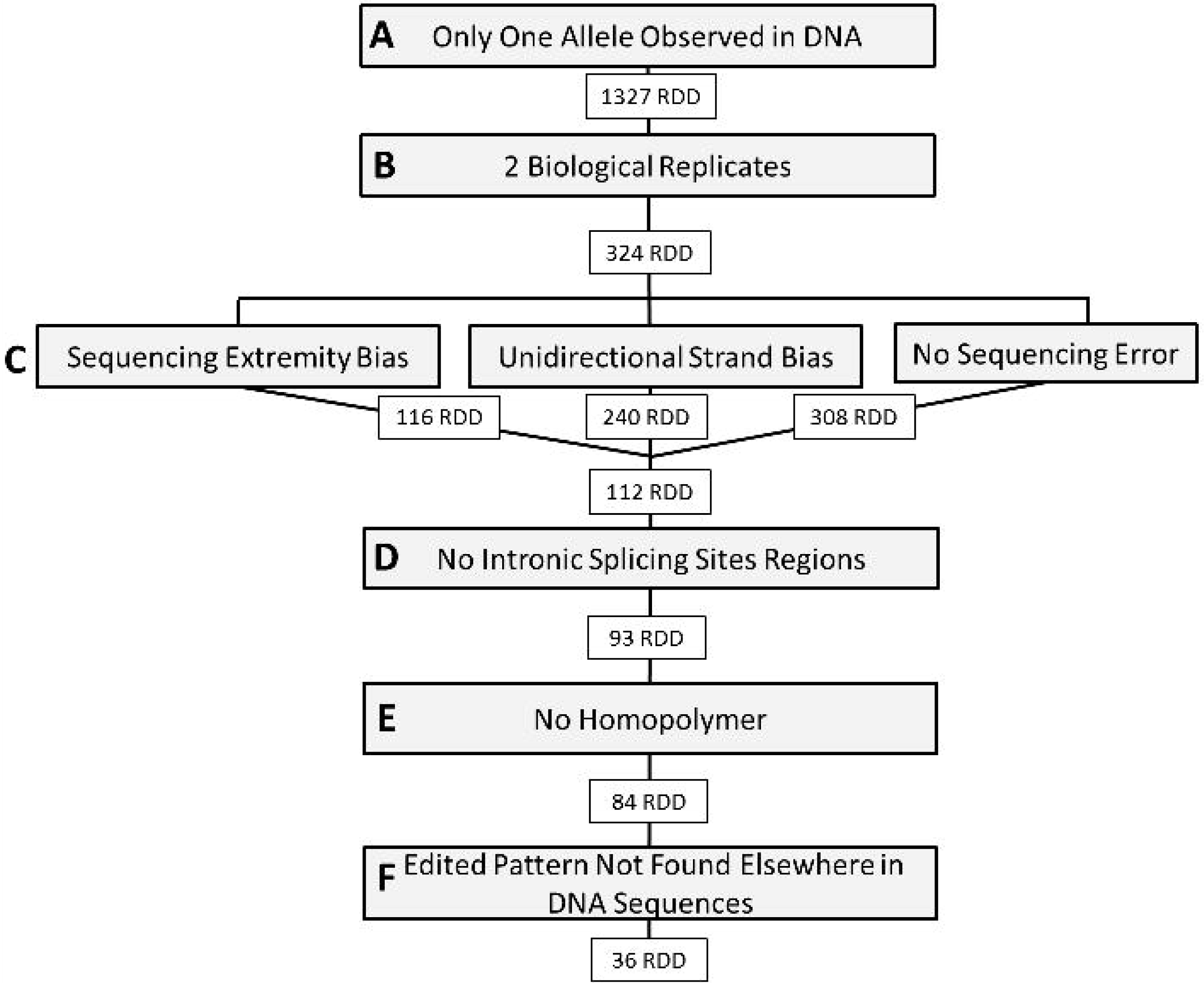
Number of RDD candidates obtained after each filter

It has previously been shown that polymorphisms overrepresented in read extremities are likely to be false positives [33–35]. In order to avoid this bias, we only considered RDD sites in which the RDD allele was, in median, not in the 10% extremities of reads overlapping them (Figure 1C). Two additional filters related to sequencing were applied: we removed candidates with an over-representation of one allele on one strand and discarded positions where more than one alternative nucleotide was found in proportions superior to 5% (Figure 1C). A total of 112 RDD sites passed all filters. We then removed candidates in splicing sites from non-coding regions (Figure 1D), and filtered for regions containing homopolymers (Figure 1E). A total of 84 candidates remained. We applied a last filter by removing candidates harboring the “edited” pattern in the genomic DNA reads (Figure 1F). The goal was here to take into account putative candidate regions for which the corresponding DNA reads were present, but unmapped or not mapped to the same position as the “edited” RNA reads.

At the end of the analysis, we found 36 reliable RDD candidates (Table 2). A total of 17 chicken genes are potentially impacted by these RDD sites, knowing that one site can be associated with several genes and that we are probably missing non-annotated genes for candidates highlighted in intergenic regions. Interestingly, many of these candidates were organized in clusters, the 36 positions corresponding to 20 different genomic regions (Table 2). A total of 7 clusters, in 5 annotated genes and in 2 intergenic regions, could be counted up, encompassing 12 to 1439 bp. The distance between 2 clustered RDDs ranged from 3 to 807 bp, for a number of detected sites comprised between 2 and 5.

**Table 2.**
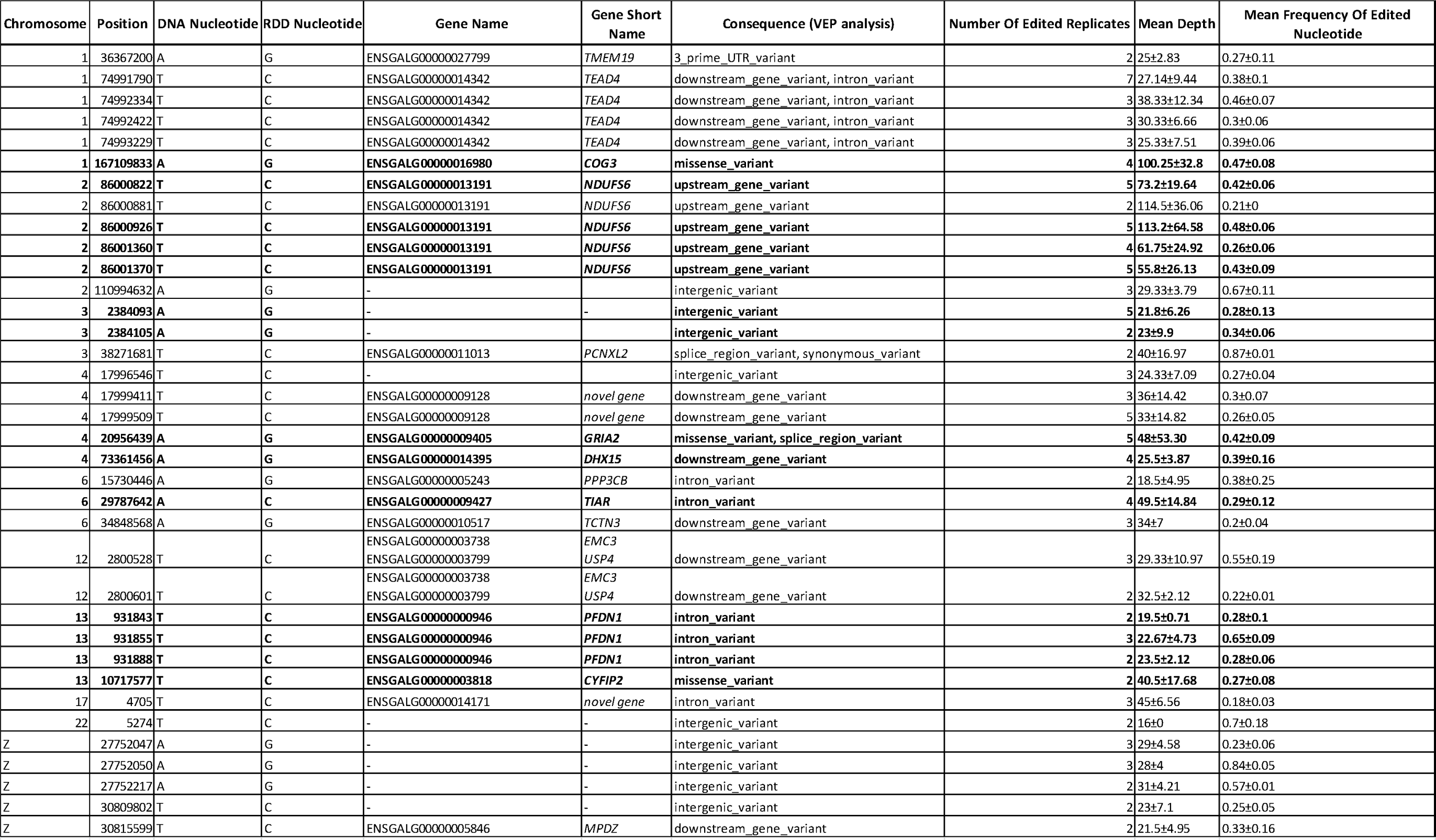
RDD candidates after filtering steps. In bold, candidates tested for validation

### RDD types

We distinguished canonical RDD (A-to-G and C-to-T) from non-canonical RDD (other base changes). As the sequencing process was not strand-specific, the complement bases of canonical changes were also considered as canonical (*i.e.* T-to-C and G-to-A).

When comparing our datasets before and after filtering, we observed a clear enrichment in canonical changes throughout successive filters, which was quite reassuring in terms of results accuracy (Figure 2). Before filtering, all possible base changes were represented, at a frequency ranging from 5 % to 20% (Figure 2). Altogether, canonical base changes represented 50% of RDD candidates. After filtering, canonical base changes represented all modifications except one, at position chr6: 29787642. This non-canonical A-to-C position seemed to be the result of a misalignment involving an alternative splice-site. This position was selected for pyrosequencing validations.

**Figure 2.**
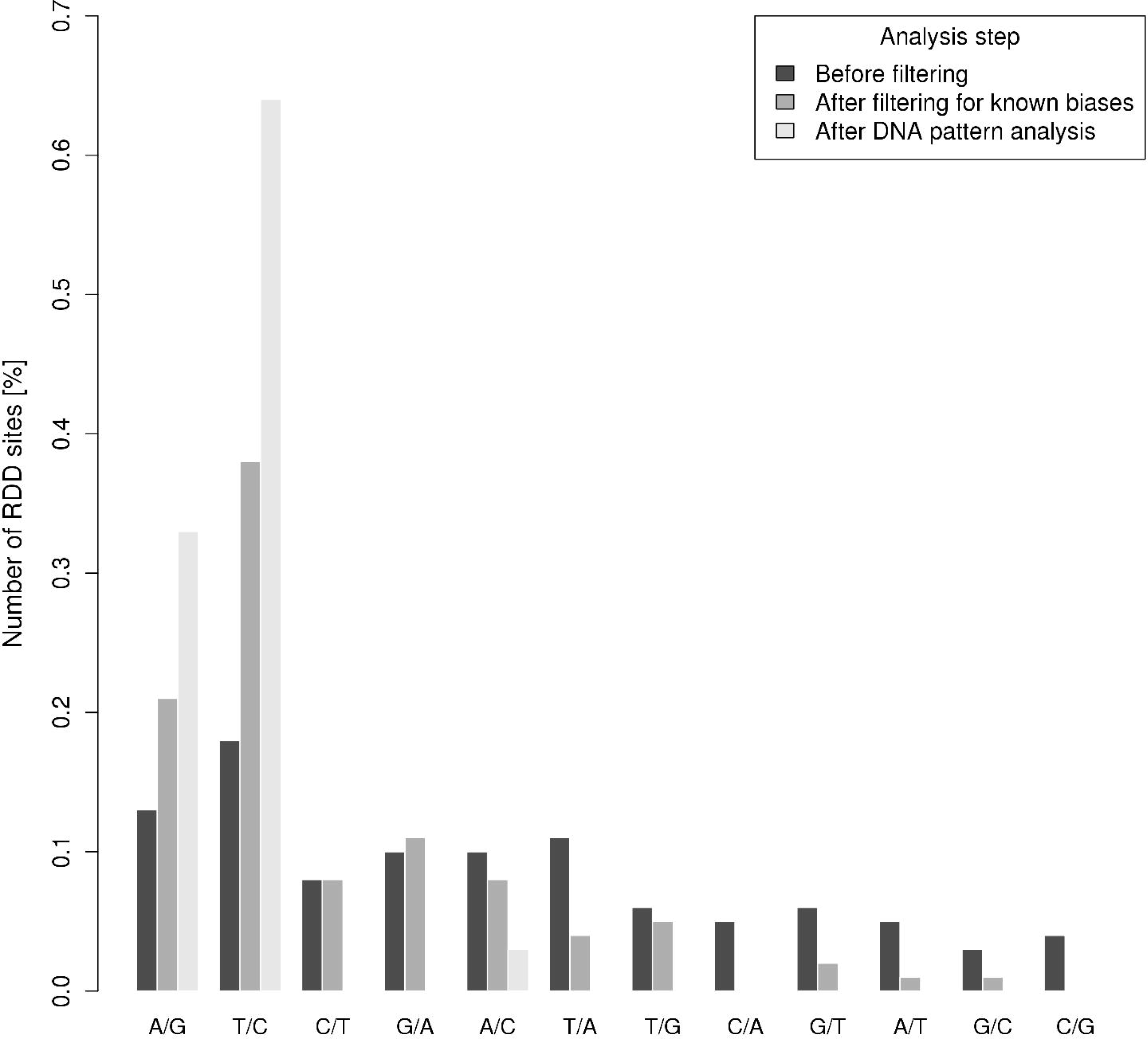
Proportion of base changes of RDD candidates before/after filters.

Among the canonical modifications, we found only A-to-G or its complement T-to-C modifications, and no C-to-T conversion.

We then characterized the RDD candidates with regards to their putative functional features.

### Functional RDD and tissue expression

Three RDD sites, located on CYFIP2, GRIA2 and COG3, were potentially functional, because leading to a non-synonymous change, and thus potentially having deleterious effects on the encoded protein. Most of the remaining candidates are located in gene introns, upstream or downstream regions of genes (Table 2, Figure 3).

**Figure 3.**
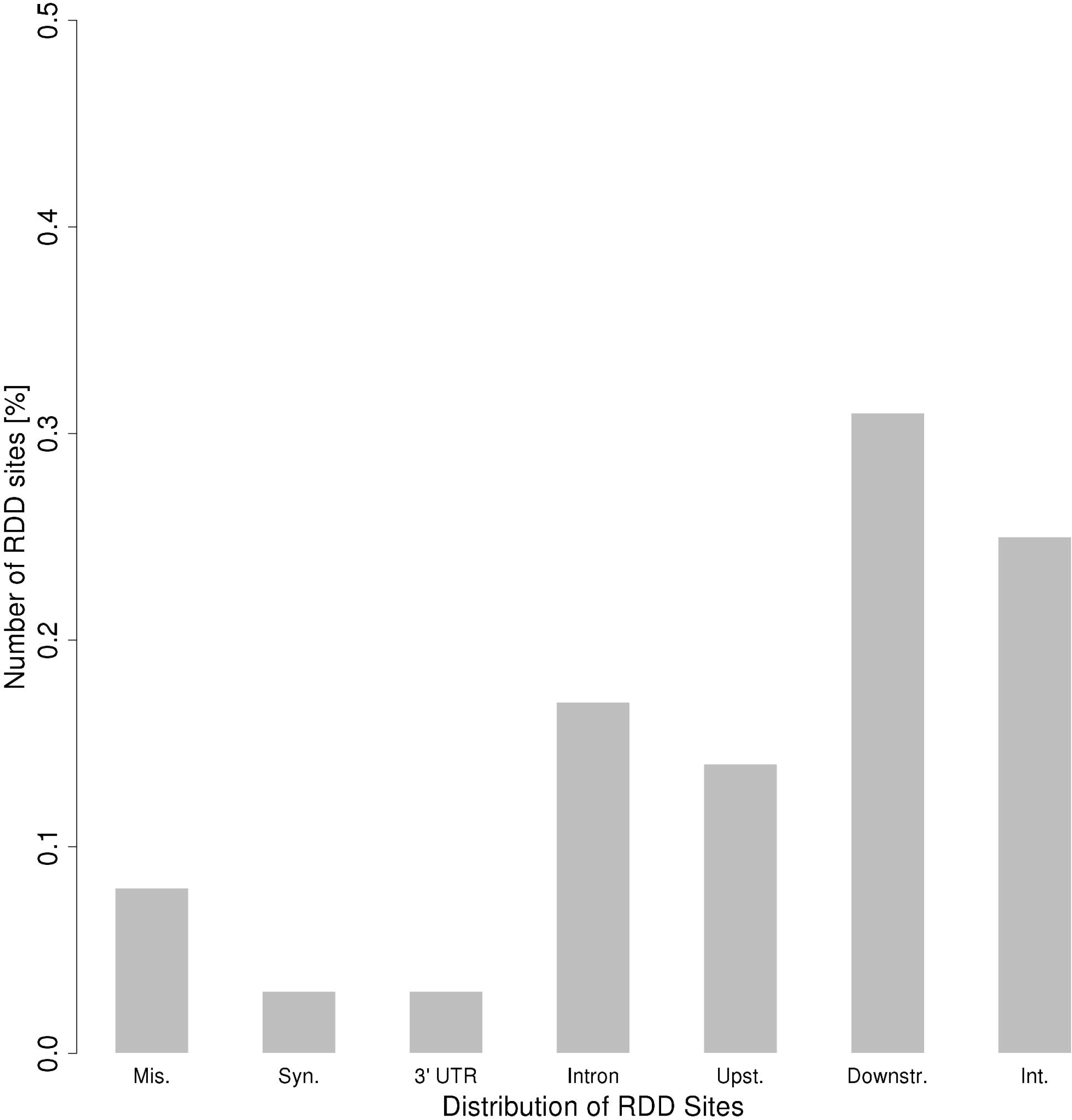
Distribution of RDD sites in genomic features.

By using 5 different *in silico* predictors of the amino-acids substitutions putative effects, we showed that none of the 3 non-synonymous substitutions was likely to be deleterious (Table 3). These substitutions were localized in highly conserved regions of the proteins (Additional file 2). A striking observation is that the K/E editing site affecting the CYFIP2 gene changes an amino-acid conserved between all examined Vertebrate species into an amino-acid which is coded without editing by the genomic sequence of Ray-finned fishes.

**Table 3.** *In silico* prediction of functional consequence on edited variants

| Variant | Software |  |  |  |  |
| --- | --- | --- | --- | --- | --- |
|  | SIFT | PolyPhen2 | Mut. Ass. | CHASM* | ProSMS |
| COG3 I632V | tolerated | benign | neutral | 0.40 (0.22) | could destabilize |
| CYFIP2 K320E | tolerated | benign | neutral | 0.32 (0.29) | no effect |
| GRIA2 R764G | tolerated | benign | medium | 0.17 (0.49) | no effect |

\* Computed using Cravat 3.0, functional score close to 1 means functional effect (score p-value)

### Characterization of candidates

We designed primers for 14 RDD candidates corresponding to 9 genomic regions, comprising missense variants, intron, upstream or downstream regions, intergenic position, and the remaining non-canonical modification. We first confirmed the homozygous status of the 14 selected RDD sites on DNA by Sanger sequencing. Their RDD status was then tested by pyrosequencing, and 13 RDD candidates were confirmed as edited loci (Figure 4). It is interesting to note that the unique site not validated by pyrosequencing corresponds to the non-canonical RDD candidate. A subset of 7 validated candidates was then tested in the other available tissues: individual heart, brain and liver tissues from three developmental times, comprising the same stage as the original HiSeq samples (day 4.5), an older embryonic stage (day 15), and an adult stage (11 months of age). Among these candidates, three are clustered on chromosome 13 (Figure5B.abc) and two are clustered on chromosome 2 (Figure 5C.ab). These positions were tested for tissue and stage effects on editing levels (Table 4). Tissue effect and stage effect were significant for all candidates (p-value≤0.05), and an interaction between tissue and stage was also observed for all but one candidates. There was a clear effect of both tissue and stage on the editing level. Interestingly, for 5 candidates out of 7, there was a continuous increase in editing level with age, from about 50% to more than 80%, independently of the tested tissue (Figure 5). In both clustered regions (Figure 5BC), all candidates harbored the same profile and only differed by their editing level. For one candidate, chr1: 167109833 (Figure 5Aa), the editing level was increasing during embryonic development and was less important in adult stage. On chr13: 10717577 (Figure5A.b), editing was mainly present in brain, with a level increase with time, and really low in other tissues. Interestingly, the editing level was tissue-specific, at every developmental time point, increasing for most of the candidates from liver to brain (Figure 5).

**Figure 4.**
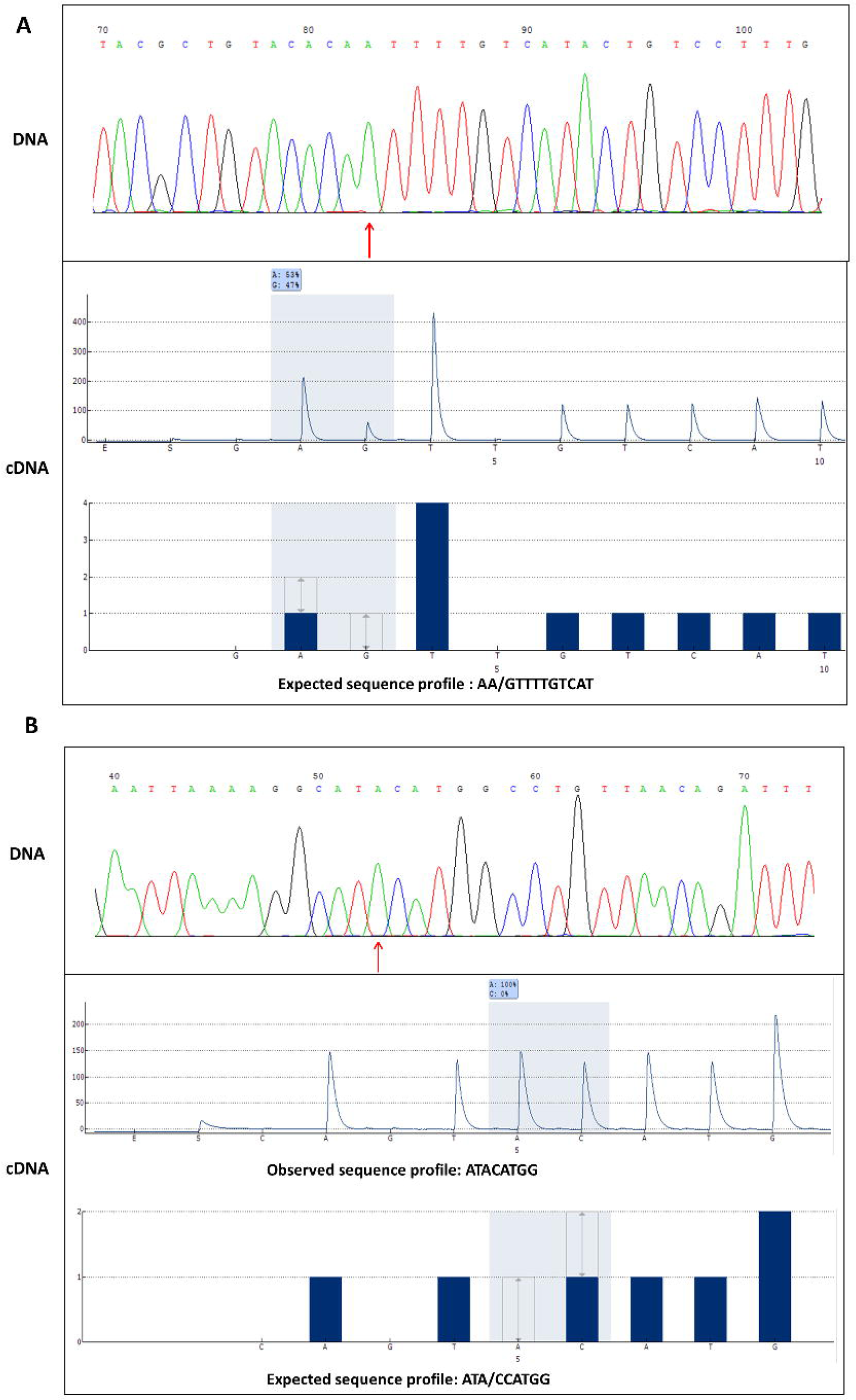
Validation of candidates by Sanger sequencing (DNA) (red arrow) and pyrosequencing (cDNA) (grey). A: Example of a canonical RDD (T-to-C) at position chr2: 86000926. The sequence is in reverse-complement. The RDD status is confirmed by pyrosequencing (A: 53% - G: 47%) **B:** Example of a non-canonical RDD (A-to-C) at position chr6: 29787642. The alternative nucleotide is not detected (A:100% - C: 0%).

**Figure 5.**
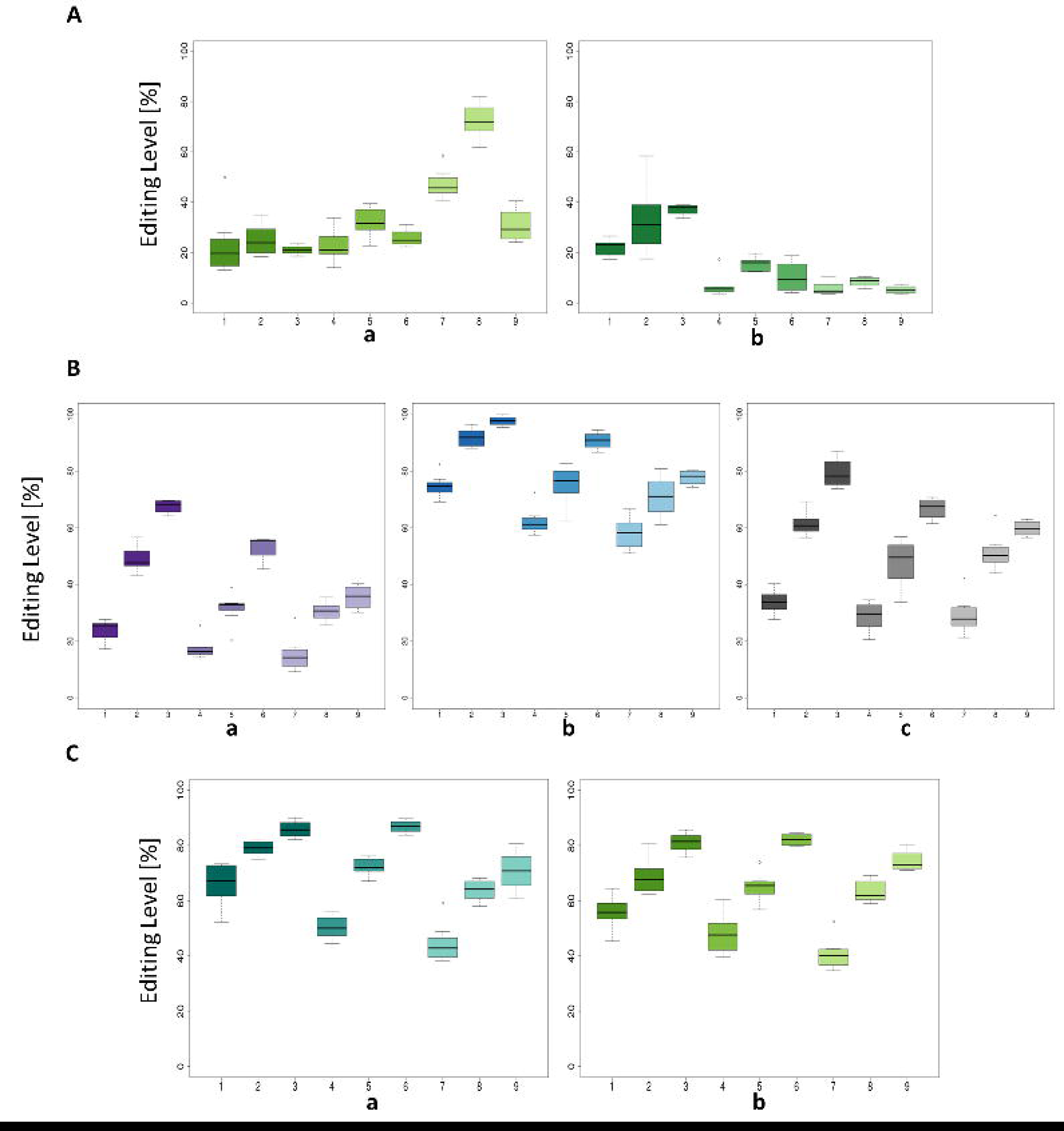
Editing levels observed across tissues and time. A: 2 selected candidates (a: chr1: 167109833; b: chr13: 10717577). B: Cluster 1 candidates (a: chr13: 931 843; b: chr13: 931855; c: chr13: 931888). C: Cluster 2 candidates (b: chr2: 86000926; c: chr2: 86001370). On abscissa axis: 1: Embryo stage 4.5 days–Brain, 2: Embryo stage 15 days–Brain, 3: Adult 11 months–Brain, 4: Embryo stage 4.5 days–Heart, 5: Embryo stage 15 days–Heart, 6: Adult 11 months–Heart, 7: Embryo stage 4.5 days–Liver, 8: Embryo stage 15 days–Liver, 9: Adult 11 months–Liver.

**Table 4.** P-values from an analysis of variance for tissue and stage effect on editing frequency. Positions of candidates tested on multiple tissues by pyrosequencing are shown.

| Chromosome | Position | Tissue effect | Stage effect | Tissus/Stage Interaction |
| --- | --- | --- | --- | --- |
| 1 | 167 109 833 | 1.29E-19 | 1.63E-09 | 1.35E-07 |
| 2 | 86 000 926 | 1.63E-14 | 1.13E-21 | 4.79E-03 |
| 2 | 86 001 370 | 8.17E-05 | 2.39E-20 | 9.94E-02 |
| 13 | 10 717 577 | 2.08E-17 | 1.11E-04 | 2.32E-02 |
| 13 | 931 843 | 1.44E-16 | 1.67E-25 | 1.45E-06 |
| 13 | 931 855 | 4.02E-15 | 1.67E-17 | 0.14 |
| 13 | 931 888 | 4.57E-08 | 4.77E-24 | 0.02 |

## DISCUSSION

Among animals, RNA editing is well described in mammals, but similar studies were lacking in other vertebrates like chicken. The goals of this study were to screen the entire chicken transcriptome for editing sites, and to characterize this phenomenon at different stages of development and tissues to extend the analysis of its conservation among vertebrates.

To do so, we used DNA-Seq and RNA-Seq technologies, allowing us to screen the whole chicken genome for such events. This approach was used recently in several species to detect RDD [12–15, 26, 27, 36–38]. While a large number of new RDD sites was first described using this approach, in particular in humans with more than 10,000 sites observed [37], these results were then contested [28, 29], showing that RNA editing would be a limited process when taking into account possible high-throughput sequencing technologies biases. Later studies confirmed this questioning by finding much less RDD sites when stringently filtering the dataset [15, 29]. For example, Pickrell and colleagues demonstrated that up to 94% of the 10,210 edited sites highlighted by Li and collaborators [37] were likely to be false positive.

We carefully looked at common analysis pitfalls when detecting RDD sites in our dataset. First, we applied a stringent filter by taking into account only RDD sites observed in at least 2 biological replicates. This filter ensures to keep true biological phenomena, and to remove candidates due to individual-dependent artefacts (as specific sequencing errors or somatic mutations putatively not seen in DNA). As there is an over-representation of mis-called SNP in read extremities [33–35], each RDD site with a biased distribution of the alternative nucleotide towards the extremities of the reads was discarded from the analyses. In accordance with previous studies [28], we chose to consider only the distribution of the “edited” nucleotide position, to increase the stringency of the method: for candidates with small RDD levels, when considering both nucleotides in the filter, the candidate can be falsely declared as unbiased and kept.

Only sites with a balanced proportion of alternative nucleotides in forward and reverse strands were kept. At this step of analysis, we found 112 RDD candidates (Figure 1ABC). Another study filtering its datasets for the same biases, with a more stringent filter concerning the number of biological replicates (at least two thirds of replicates detected as edited) [15], found between 128 (in mouse liver) and 447 (in mouse adipose) RDD candidates at the same filtering step, *i.e.* more candidates than our results while their study was limited to the exome. It could constitute an argument in favor of the scarcity of RNA editing in chicken. We chose to be really stringent by keeping only RDD positions with a total absence of the alternative nucleotide on DNA. It appears that the SAMtools mpileup SNP detection software declared homozygous DNA positions where we could find the alternative nucleotide harbored by several reads. We are aware of the possible loss of real candidates, but the aim was here to maximize the reliability of the results. The final step, eliminating candidates for which the edited pattern was found in the DNA reads, removed 48 candidates, even if they passed our stringent filters. At the end of the filtering steps, we kept less than 10% of putative RDD sites detected at the beginning of our study, which is similar to results obtained in recent studies, taking into account biases linked to high-throughtput sequencing [15, 17].

Compared with previous studies, a distinguishable feature of our analysis is the search of the “edited” pattern in the DNA reads of candidates highlighted through RNA-Seq. In several cases, while the “edited” RNA reads map to a candidate region, the corresponding DNA reads do not map to the chicken genome: the RNA read can be mapped to a paralogous region due to the splitting of introns, while the original region is either absent from the genome or is carrying too many mismatches between our individuals and the reference sequence. This can be explained by an incomplete genome assembly and / or several regions with assembly errors. Indeed, the chicken genome assembly is still incomplete, especially regarding microchromosomes [39, 40]. The false RDD status of many candidates due to DNA polymorphism in paralogous regions has already been highlighted [14]. A similar observation has been made by Piskol *et al* [30], leading to the conclusion that non-canonical editing site are likely to be false positive RDDs. Previous studies successfully detected a high number of edited sites in human [13, 36], mainly in Alu sequences. These human repeated sequences clearly show a high propensity to harbor A-to-I editing sites, which is a strong argument in favor of their true existence, and the human genome assembly is more complete than the chicken one (but with much more repeated regions). But given the results obtained in our study, we confirm that a part of the putative edited sites obtained from RNA-Seq data may be removed when taking into account not only the DNA sequence from the same sample or the biases previously underlined as the position in the reads or the strand bias, but also the putative alignment problems due to the genome assembly or to the discontinuous nature of RNA-Seq data.

At the end of the filtering steps, the number of RDD candidates was considerably reduced. Our filters were quite stringent, and we may have missed a few real positions. But as we still detected a false positive candidate through an experimental validation, this high stringency was surely appropriate. We could also have missed true candidates because of the alignment stringency: if some regions are extensively edited, the resulting RNA sequence becomes really different from its DNA matrix. As a consequence, reads sequenced from edited RNA carry too many mismatches to be kept in the alignment [9, 41], except when using appropriate computational methods [42]. But given the small extent of RDD in chicken, this hypothesis is not likely.

Nevertheless, this low number of candidates tends to suggest that, as in other non-primate animals, RNA editing is a limited event in chicken [43].

The proportion of canonical RDD changes increased across filters, which is reassuring about the reliability of the pipeline: only one non-canonical change could be observed, shown not to be a true conversion, due to misalignments along an alternative splice site. The status of this false candidate was confirmed by pyrosequencing.

Interestingly, no C-to-T conversion was observed in our dataset. In particular, confirming previous studies on RNA editing in chicken, we could not find any editing in APOB transcripts [19]. This is in accordance with the missing of APOBEC1 from the chicken genome, as this enzyme seems to be required for C-to-U APOB RNA editing in vertebrates [5].

After the detection of RDD sites in the chicken transcriptome, our aim was to further characterize several interesting edited positions.

All the tested candidates were shared between the studied tissues in our analysis, with one candidate presenting a very low level of editing in the liver, whatever the stage (chr13: 10717577, Figure 5Ab). But the edition level varies between the analyzed tissues and ages, confirming in our chicken model that RNA editing varies across times and tissues.

Interestingly, RNA editing levels change over time. As generally observed in mammals, with few exceptions [44–47], the A-to-I editing level increased during development, as it is the case for 6 out of 7 tested candidates. Even if the tissue and stage specificity of edition is clear in candidates tested from cluster regions (Figure5BC), it is even more pronounced for tested candidates highlighted separately (Figure5A). These time- and tissue-specific phenomena are not only due to the level of expression of ADARs [46, 48] and more work is needed to decipher the spatio-temporal regulation of RNA editing. The low level of edition at embryonic stages in almost all the tested candidates could be explained by a putative importance for adequate embryologic development, as it was hypothesized for the GRIA2 Q/R site in mammals [46], even if it has to be confirmed.

As previously highlighted, our results confirm that editing at a particular position often comes with editing sites nearby, but no clear functional explanation has been proposed yet [46]. The regional sequence composition and RNA molecule tertiary structure seem to be involved in these clustered editing sites [49, 50].

One interesting result is that only a few candidates were directly affecting the protein sequence by changing an amino-acid. It has been shown that RNA editing can impact protein function, like modifying ions channels in some tissues [51, 52], or impacting the ligand-binding affinity [53]. Nevertheless, our results show that RNA editing in chicken is more frequently silent, as already observed [54]. More studies should be performed to confirm these results. But a significant number of candidates are located in non-coding parts of the chicken genome, at least given the current state of the annotation. As in a previous study in human Alu regions [49], we observed a high number of edited sites in introns. Even if our data come from polyA+ RNAs, these sites may correspond to editing in pre-mRNAs. But they may be part of non-coding RNAs too, where editing has been discovered in several species, and the biological significance of which is still largely unknown [54, 55].

We highlighted 3 candidates that were previously described as edited in mammals, one K/E substitution already observed in the CYPIF2 gene [21], one I/V conversion located in the COG3 gene [16, 17, 56] - previously described in human, mouse and rat - and the R/G site in the GRIA2 gene [57], which means that these editing events are not restricted to mammals and appeared before the Sauropsid-Synapsid divergence. Possible implications of an altered editing efficiency at the R/G site in GRIA2 in mental disorders in human and mouse were recently observed [58]. Concerning the COG3 editing site, no functional implication is documented at this time, but as underlined in another study [16], the conservation of this site both in mammals and in a broader way in vertebrates implies a putative functional role. Similarly, the functional significance of the conserved CYFIP2 K/E editing, which is higher in brain than in other tissues in human, is not known, but may be implicated in apoptosis [45].

The editing sites are located in highly conserved regions between vertebrates (Additional file 2). Interestingly, the modification observed in the CYFIP2 gene results in a conversion from a Glutamic acid to a Lysine. This amino-acid is only present in the Ray-finned fishes, and shared by all of them (http://www.ensembl.org/index.html). The other species for which the homologous CYFIP2 sequence is available are all K-coding at this position, which asks the question of the functionality of this E residue, only present in fishes as a chromosomal codon, but resulting from an editing phenomenon in several Vertebrate species, including chicken.

The very small number of conserved edited sites between species has already been underlined [59] and may be the signature of their functional importance.

## CONCLUSIONS

This study constitutes, to our knowledge, the first whole genome screening of RNA editing in chicken. By using a stringent pipeline, we focused on really reliable RNA editing events and thus removed most putative false positives, a big pitfall in RNA editing discovery through high-throughput sequencing. Our pipeline predicts reliable RNA editing site; most of the tested sites are confirmed through an independent validation method, avoiding biases encountered when using NGS data. RNA editing seems to be a very limited phenomenon in chicken, at least in whole embryo at 4.5 days of age, as attested by a whole genome screening through RNA-Seq. This whole genome analysis shows that the A-to-I editing mechanism may be the only one present in chicken. Several edited loci are conserved between chicken and other vertebrate species, including human, which indicates that, while RNA editing arose long ago in the evolution, some particular nucleotides from a few genes are subject to RNA editing. This conservation is probably linked to the molecular mechanisms involved, but more deeply questions the functionality of editing at these specific loci. Even if the spreading of RNA editing is more and more characterized, a huge effort to discover the putative functionality of this phenomenon is still needed.

## METHODS

### Tissues dataset

The material used in this study for the embryo sequences dataset was previously described [60] [SRA study accession number: SRP033603]. Briefly, two chicken lines were crossed, Line 6 [61] and Line R^−^ [62]. Chickens were bred at INRA, UE1295 Pôle d’Expérimentation Avicole de Tours, F-37380 Nouzilly in accordance with European Union Guidelines for animal care. Twelve F1 were produced from 2 families: 8 embryos (embryonic day 4.5) and 4 adults from the same batch. Embryos were kept as a whole, while 3 adult tissues were harvested: brain, heart, and liver. Additional embryos were produced at embryonic days 4.5 (n=8) and 15 (n=8), from a cross between the same lines, and 3 embryonic tissues were harvested: brain, heart and liver. Genomic DNA and total RNA were concurrently extracted from the same samples of crushed whole embryos or individual tissues (AllPrep DNA/RNA Mini Kit, Qiagen). RNA quality was measured by a BioAnalyzer (Agilent); all samples had a RIN (RNA Integrity Number) ≥ 9.9.

### Sequencing

#### RNA sequencing

Libraries with a mean insert size of 200bp were prepared following Illumina instruction for RNA-Seq analysis, by selecting polyA+ fragments (TruSeq RNA Sample Prep Kit) from each sample. Samples were tagged to allow subsequent identification, amplified by PCR and quantified by qPCR (Agilent QPCR Library Quantification Kit).

A total of 8 embyo libraries were sequenced (paired-ends, 100 bp) in triplicate on an Illumina HiSeq 2000 sequencer (Illumina, TruSeq PE Cluster Kit v3, cBot and TruSeq SBS Kit v3) by randomizing their position in 6 different sequencing lanes.

#### DNA sequencing

DNA from 8 embryos was sequenced on 5 lanes of Illumina HiSeq 2000. Library preparation (mean insert size 328bp), DNA quantification and sequencing (paired-ends, 100bp) were performed according to the manufacturer instructions (TruSeq DNA Sample Prep Kit Illumina, Agilent QPCR Library Quantification Kit, TruSeq PE Cluster Kit v3 cBot TruSeq SBS Kit v3).

### Computational analyses

When not specified, analyses were performed with homemade Perl and R scripts.

#### Genomic sequences analyses

Sequences were aligned to the current chicken genome assembly (Gallus gallus 4) using the BWA program version 0.7.0, option aln [63]. Sequences were then filtered on mapping quality (MAPQ SAMtools rmdup command was used to remove possible PCR duplicates.

#### PolyA RNA sequences analysis

Sequences were aligned with Tophat software version 2.0.5 on the chicken reference genome Galgal4 as described in [60].

Sequences mapping uniquely on the reference genome, without PCR duplicates and with a minimum mapping quality of 30, were selected.

#### Identification of RNA/DNA differences

Sequences were locally realigned and recalibrated before SNP detection with GATK software version 1.6.11 and BamUtil (bam recab command).

SAMtools software version 0.1.19 was used with mpileup utility to detect SNPs between DNA and RNA samples from each individual. We set a maximum coverage of 10,000 for each calling to take into account as many reads as possible in the calling. SNPs were detected independently on each biological replicate.

#### Editing detection

SNPs were analyzed from VCF files obtained from SAMtools mpileup detection. For each biological replicate, only variations where DNA was homozygous either for the reference allele or for the alternative allele, and where RNA was heterozygous, were kept.

Several successive filters were applied to consider a position as a putatively RDD site. We first only considered positions with a sufficient depth, keeping only candidates presenting a minimum of 15 reads both in DNA and RNA alignments. To increase the likelihood of a site to be a true RDD position by avoiding a sample artefact, we set to 2 the number of biological replicates that must carry the same modification.

We then applied several filters inherent to technical bias due to high-throughput sequencing. Positional bias was checked and all RDD candidates for which the median position of the “edited” allele among reads overlapping them was in the 10 first or 10 last bases were discarded. The strand bias was also considered; to be kept, a RDD candidate must present a proportion of edited allele on the forward strand really close to its proportion on reverse strand (delta ≤0.5).

We checked the biallelical status of each selected candidate, a third allele being detected in less than 5% of cases being considered as a sequencing error.

An additional filter was applied to ensure that the alternative nucleotide frequency on DNA was null. The functional consequence of each RDD in each transcript was predicted using the Ensembl Variant Effect Predictor (VEP) version 71 [64]. Non-coding splicing site regions were removed to take into account putative misaligned reads at these sites [31]. Then, positions belonging to homopolymers (n≥5) were removed because they may generate false positive candidates [13].

The chicken genome assembly still lacks several assembled regions, due to sequence assembly errors or missing fragments. A fragment detected as uniquely mapped may thus be present, with several polymorphisms, at genomic regions absent from the reference sequence, but present in the DNA reads from our samples. Therefore, a last filter was performed by searching the “editing site” (40 bp surrounding the candidate locus) in the DNA reads from samples thought to be edited. This pattern was searched with fuzznuc [65].

## Validation assays and editing characterization

### Sanger sequencing

We first checked the homozygous status of RDD sites by Sanger sequencing on DNA. The 8 biological replicates were tested. Primers were designed using PyroMark Assay Design software to allow further cDNA pyrosequencing (Additional file 1).

### Pyrosequencing

RDD sites were tested on a Qiagen PyroMark Q24 sequencer. Primers were designed with PyroMark Assay Design software (Additional file 1). PCR products were made using PyroMark PCR Kit (Qiagen). We performed the analyses through the PyroMark Q24 1.0.10 software with default analysis parameters.

Tissue and stage effects on the editing level were tested through an analysis of variance in a model taking into account tissues, stages and the interaction between tissues and stages for each tested candidate.

### In Silico prediction of protein structure and function

To predict the putative effect of the editing conversions on protein structure and function, we used several bioinformatic tools: SIFT is based on sequence homology and the physical properties of amino acids (http://sift.bii.a-star.edu.sg/); PolyPhen2 uses physical and comparative considerations (http://genetics.bwh.harvard.edu/pph2/); MutationAssessor is based on evolutionary conservation of the affected amino acid in protein homologs (http://mutationassessor.org/); CHASM (computed using CRAVAT 3.0: http://www.cravat.us/) is based on the probability that a modification gives the cells a selective survival advantage; ProSMS predicts protein stability changes due to single amino acid modifications (http://babel.ucmp.umu.se/prosms/).

## ACKNOWLEDGMENTS

We thank the entire staff of the PEAT experimental unit for their excellent animal care, and Juliette Riquet, Julie Demars, Gwenola Tosser, Annie Robic and Bertrand Servin for helpful discussions about the results presented in this study. Sequencing was performed at GeT-PlaGe Genotoul platform.

*This paper is dedicated to the memory of André Bordas*.

## ADDITIONAL FILES

**Additional file 1 (Fresard_Table S1.xls): Sequencing primers.** [Btn] Biotin on 5' end for pyrosequencing only.

**Additional file 2 (Fresard_Figure S1.tiff): Alignment of protein sequence from different species.**

Multi-species alignments were performed through the Muscle program in the PhyleasProg pipeline (phyleasprog.inra.fr), from reference protein sequences of fully sequenced genomes from Ensembl (www.ensembl.org).

The red arrows show the amino acid affected by the editing conversion. The overall conservation between all species is depicted under each multi-alignment. A. COG3 (I-->V) B. GRIA2 (R-->G) C. CYFIP2 (K-->E).

